# *Plasmodium falciparum* mitochondrial complex III, the target of atovaquone, is essential for progression to the transmissible sexual stages

**DOI:** 10.1101/2024.01.09.574740

**Authors:** Pradeep Kumar Sheokand, Alexander Mühleip, Lilach Sheiner

## Abstract

The Plasmodium mitochondrial electron transport chain (mETC) is responsible for essential metabolic pathways such as de novo pyrimidine synthesis and ATP synthesis. The mETC complex III (cytochrome bc1 complex) is responsible for transferring electrons from ubiquinol to cytochrome c and generating a proton gradient across the inner mitochondrial membrane, which is necessary for the function of ATP synthase. Recent studies revealed that the composition of Plasmodium complex III is divergent from human, highlighting its suitability as a target for specific inhibition. Indeed, complex III is the target of the clinically used anti-malarial atovaquone and of several inhibitors undergoing pre-clinical trials, yet its role in parasite biology have not been thoroughly studied. We provide evidence that the universally conserved subunit, PfRieske, and the new parasite subunit, PfC3AP2, are part of Plasmodium falciparum complex III (PfCIII), with the latter providing support for the prediction of its divergent composition. Using inducible depletion, we show that PfRieske, and therefore PfCIII as a whole, is essential for asexual blood stage parasite survival, in line with previous observations. We further found that depletion of PfCIII results in gametocyte maturation defect. These phenotypes are linked to defects in mitochondrial functions upon PfRieske depletion, including increased sensitivity to mETC inhibitors in asexual stages and decreased cristae abundance alongside abnormal mitochondrial morphology in gametocytes. This is the first study which explores the direct role of the PfCIII in gametogenesis via genetic disruption, paving the way for a better understanding of the role of mETC in the complex life cycle of these important parasites and providing further support for the focus of antimalarial drug development on this pathway.

## Introduction

Malaria is a deadly disease which causes significant morbidity and mortality around the globe with 249 million cases and ∼608,000 deaths reported in 2022 alone[1]. It is caused by mosquito-transmitted parasites of the *Plasmodium* genus, with *Plasmodium falciparum* being responsible for most fatal cases in human. The parasites have a complex life cycle with several life stages parasitising different tissues in the host and in the vector [2]. In the blood of the human host, the parasite life stages can be divided to the asexual stage, responsible for the disease symptoms, and the sexual stage (gametocyte) responsible for the disease transmission to the mosquito vector. Mosquito can take up the gametocytes during a bloodmeal thus initiating transmission. Drugs targeting both these stages exist and are use as one of the strategies for malaria control [3, 4]. The rapid emergence of drug resistance however threatens the current first-line therapies and highlights a need to develop new compounds [5]. A better understanding of pathways that are known to be drug targets may contribute to this goal.

One of these pathways is the *Plasmodium* mitochondrial electron transport chain (mETC). This pathway harnesses electrons from different metabolic pathways via a series of dehydrogenases, including malate quinone oxidoreductase (MQO), dihydroorotate dehydrogenase (DHODH), succinate dehydrogenase (complex II) and a type II NADH dehydrogenase (NDH2). The electrons are transferred between complexes of the mETC via the mobile carriers ubiquinone, which transfers electrons to complex III (also known as the cytochrome bc1 complex), and cytochrome c, which transfers them to complex IV. At complex IV these electrons reduce oxygen to water. The main function of mETC mediated electron transfer in asexual parasites is the electrons harnessing necessary for the pyrimidine synthesis pathway [6]. Additionally, the movement of the electrons through the mETC complexes III and IV is coupled to the pumping of protons which generates a proton gradient across the inner mitochondrial membrane. The proton gradient is utilised by ATP synthase (complex V) to generate ATP. This process is collectively named oxidative phosphorylation (OXPHOS). While not essential for asexual blood stages (which generate most of its ATP via cytoplasmic glycolysis) ATP synthase becomes essential at the mosquito stage [7]. Therefore, the role of mETC, is essential in both critical stages of the parasite life cycle.

The *Plasmodium falciparum* mETC complex III (PfCIII) is a well-established drug target for the clinically used drug, Atovaquone (ATQ) [8-10] and for inhibitors undergoing pre-clinical trials such as the series of endochin-like quinolones (ELQs) [11-13]. In mammalians three subunits compose the catalytic core of complex III: 1) the mitochondrially encoded cytochrome b (cytb), which spans the inner mitochondrial membrane and thus connects the two substrate binding sites named Qo and Qi; 2) cytochrome c1 (cytC1) and 3) Rieske. The last two together mediate the transfer of electrons between the two mobile carriers. The human complex III consists of eight further non-catalytic subunits [14]. PfCIII is proposed to consist of 12 subunits, which include homologs of cytb, cytC1 and Rieske, but not all eight non-catalytic subunits found in human have homologs [15]. Moreover, additional three apicomplexan specific subunits (PfC3AP1, PfC3AP2, PfC3AP3) are proposed to be part of PfCIII [15]. Despite its divergence and importance as a drug target, the composition and function of PfCIII is not fully understood. In this study we provide support for the previously proposed divergent PfCIII composition through co-immunoprecipitation, localization, and gel migration of endogenously tagged proteins, while focusing on one conserved (PfRieske) and one apicomplexan specific subunit (PfC3AP2). Using conditional knockdown, we demonstrate that PfRieske is required for PfCIII function, and confirm that the complex is required for asexual blood stage survival whereby PfRieske depleted parasites show a defect in pyrimidine synthesis and become hypersensitive to mETC inhibitors, in line with previous reports of PfCIII disruption [10, 16]. We further found that PfRieske depletion resulted in a defect in gametocyte stage progression which coincides with a defect in mitochondrial cristae formation. This work provides support to the predicted role of PfCIII in the mitochondria of asexual and sexual *Plasmodium* stages and highlights the pathways it contributes to.

## Results

### PfRieske and PfC3AP2 are part of Plasmodium falciparum complex III (PfCIII)

Recent studies of the apicomplexan mETC complex composition via complexome profiling in both Plasmodium falciparum and *Toxoplasma gondii* have shown that their complex III consist of 12 and 11 subunits respectively [15, 17]. Both studies further identified new proteins that have no clear homologs in the mammalian complex [15, 17]. In Plasmodium three new proteins (PfC3AP1, PfC3AP2, PfC3AP3) are annotated as putative PfCIII subunits. To provide support for this divergent subunit composition we created tagged lines of two subunits, the parasite specific PfC3AP2 and the conserved PfRieske. While engineering the tagged lines we already included also the glmS system for inducible gene depletion [18]. Thus, we generated PfC3AP2-HA-glms and PfRieske-HA-glmS transgenic parasite lines whereby the 3’ region of each gene is fused with the HA-glmS tag, and the fusion proteins are expressed under the native promoter of the respective genes (Figure 1A). The correct integrations of the HA-glmS tags in the target loci was confirmed by PCR analysis (Figure 1B) and by western blot analysis (Figure 1C). Native migration and western blot analysis, used previously to resolve mETC complexes in the related apicomplexan *T. gondii* [17], further showed that both the proteins migrated at the same size of ∼730 kDa (Figure 1D) in line with the predicted size of a PfCIII dimer based on its proposed composition [15]. This provided support that both proteins are part of the complex. Further, we performed a single immunoprecipitation experiment under native conditions with the PfRieske-HA-glmS transgenic parasite line (Figure 1E) followed by mass spectrometry analysis (Table S1). We found seven of the expected twelve subunits (Table 1) including PfC3AP2, providing support for the proposed composition and further validating that both subunits are part of PfCIII.

**Table 1.** The interacting partners of the PfRieske identified via immunoprecipitation and mass spectrometry analysis. Full MS data in Table S1.

| Expected | Detected | Mass (Da) | Accession |
| --- | --- | --- | --- |
| Cytb | Yes | 43747 | PF3D7_MITO2300 |
| Cytc1 | Yes | 46613 | PF3D7_1462700 |
| Rieske | Yes | 41354 | PF3D7_1439400 |
| QCR6 | No | 1100 | PF3D7_1426900 |
| QCR7 | Yes | 23119 | PF3D7_1012300 |
| QCR8 | No | 17000 | PF3D7_0306000 |
| QCR9 | No | 11700 | PF3D7_0622600 |
| MPPalpha | Yes | 56440 | PF3D7_0523100 |
| MPPbeta | Yes | 62189 | PF3D7_0933600 |
| C3AP1 | No | 9500 | PF3D7_0722700 |
| C3AP2 | Yes | 19557 | PF3D7_1326000 |
| C3AP3 | No | 6200 | PF3D7_0817800 |

**Figure 1.**
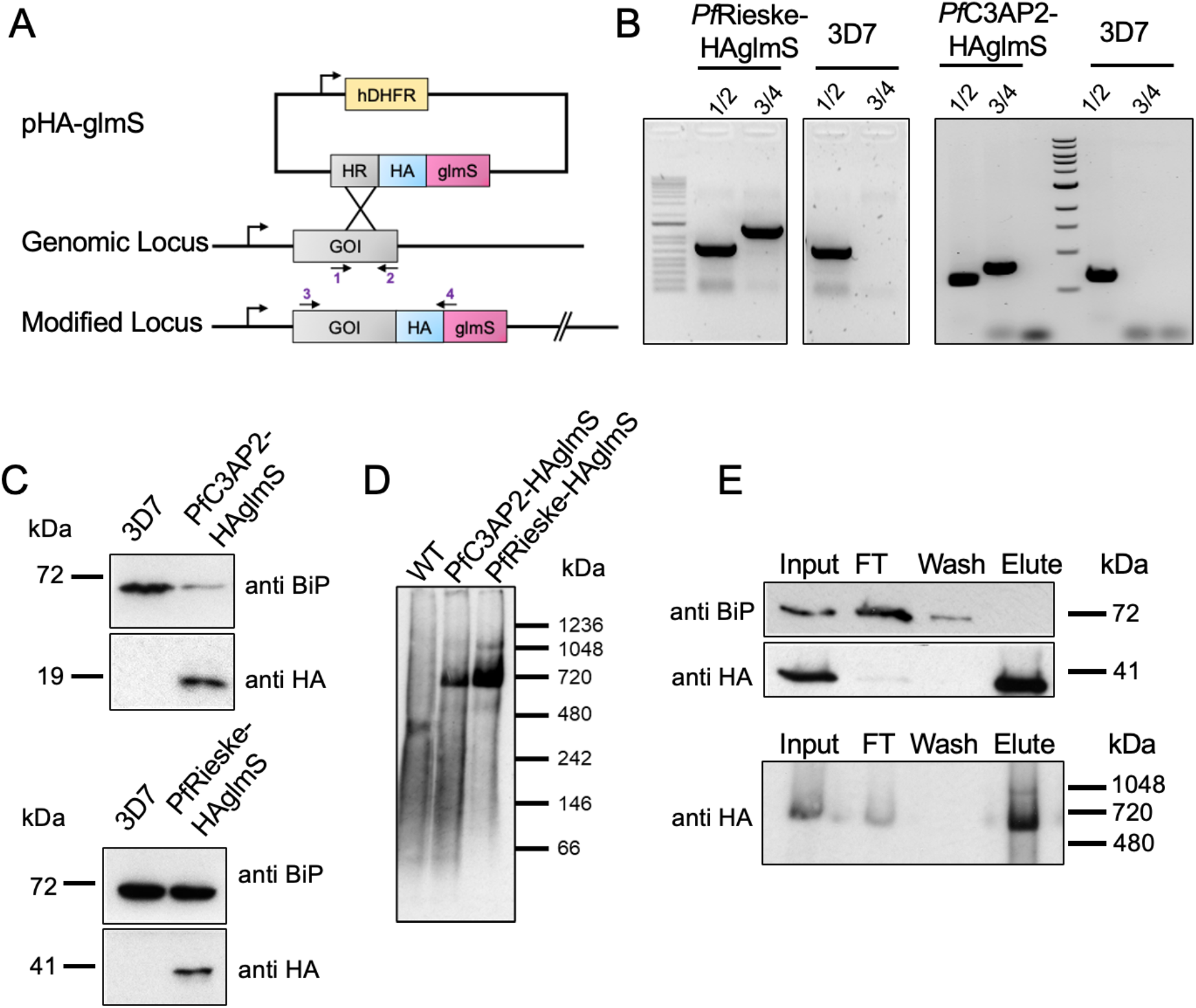
Tagging analysis and Immunoprecipitation of the PfRieske and PfC3AP2: (A) Schematics of the endogenously C terminal tagging strategy. (B) PCR analysis confirming the regulatable cassette integration using primers corresponding to the scheme in (A). (C) SDS-PAGE western blot of endogenously tagged PfRieske and PfC3AP2 using anti HA antibody and anti BiP as a loading control. (D) BN-PAGE western blot of both tagged proteins using anti HA antibody. (E) Denaturing (top panels) and native (bottom panel) PAGE and western blots of immunoprecipitation fractions from PfRieske-HA-glmS line, showing the HA enrichments and the high migrating complex found in the elute. Anti-BiP antibodies were used as fraction specificity control. Input – enriched mitochondria; FT – flow through or unbound material.

Next, we used the endogenous tags to analyse the localisation of both proteins using anti HA antibody via immunofluorescence. We observed the signal in both asexual blood stages as well as gametocytes stages (marked with the gametocyte marker Pfs25) of the parasite. The signal could not be detected in asexual ring stages, potentially due to the small size of their mitochondria reducing detection, but manifested as a tubular shape in trophozoites and became branched in the schizonts (Figure 2A, Figure 3A). In gametocytes the signal is seen to be in elongated tubular like structure (Figure 2B, Figure 3B). These observations are in line with a mitochondria shape [19, 20]. In agreement with this, both the proteins co-localise with Mitotracker Deep-Red which confirms the predicted localisation of these proteins to the mitochondria (Figure 2C-D. Figure 3C-D).

**Figure 2.**
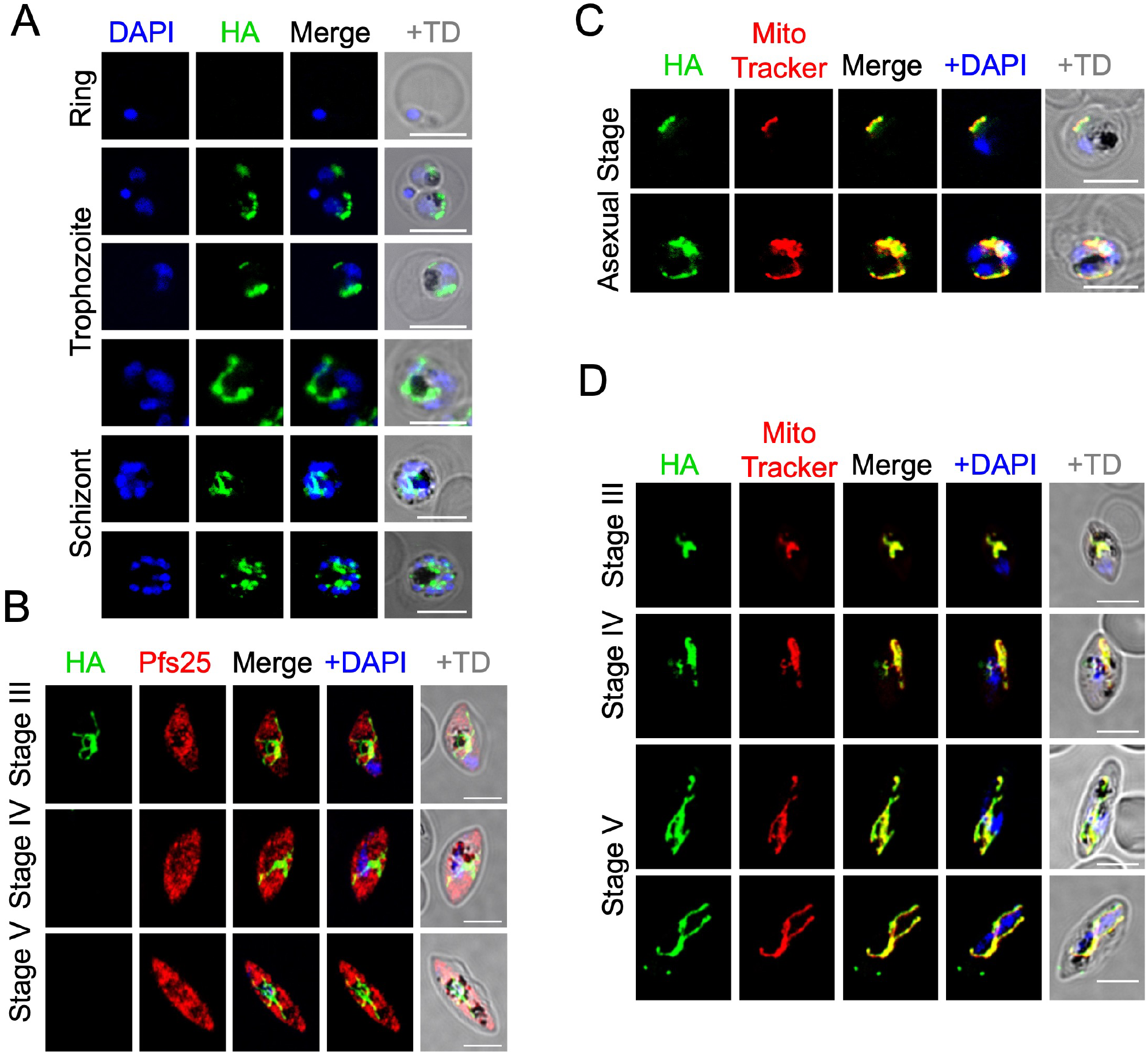
Expression and subcellular localisation of PfRieske in *P. falciparum*. (A,B) Immuno-fluorescence assay of PfRieske, visualised with anti-HA, in asexual blood stages (A) and in gametocytes (B). (C,D) Immunofluorescence assay of PfRieske visualised with anti-HA and co-stained with MitoTracker Deep-Red showing colocalization in both asexual (C) and sexual (D) blood stage parasites. Scale bar: 5μm.

**Figure 3.**
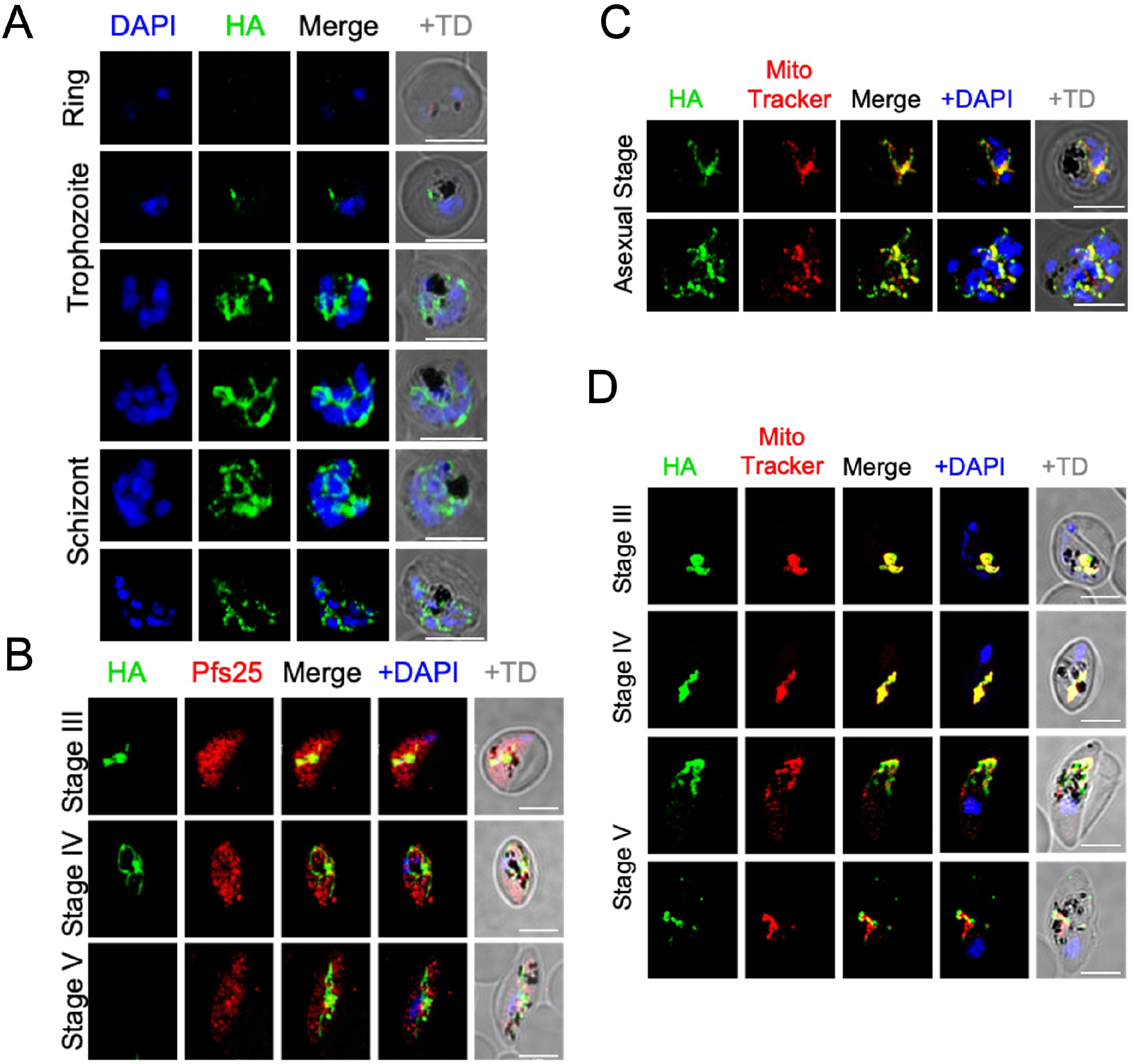
Expression and subcellular localisation of PfC3AP2 in *P. falciparum*. (A,B) Immuno-fluorescence assay of PfC3AP2, visualised with anti-HA, in asexual blood stage (A) and in gametocytes (B). (C,D) Immunofluorescence assay of PfC3AP2 visualised with anti-HA and co-stained with MitoTracker Deep-Red showing co-localisation in both asexual (C) and sexual (D) blood stage parasites. Scale bar: 5μm.

### PfRieske is essential for parasite growth

The importance of PfCIII for Plasmodium survival is well established both based on numerous inhibitor sensitivity and resistance mutant analyses [21-25], and, more recently, through the study of the subunit cytC1 [16]. However, the impact of PfCIII disruption on parasite biology has not been fully characterise. To address this gap, we focused on the two subunits we studied above and attempted their genetic disruption. We used the HA-glmS lines to induce a knockdown of each gene. Western blot analyses using the HA tag showed a significant decrease in the levels of both PfC3AP2 and PfRieske when parasites were grown in the presence glucosamine (GlcN) (Figure 4A,B). To address the effect of these knockdowns on parasite growth, we cultured the PfRieske-HA-glmS and PfC3AP2-HA-glmS parasites in presence of the GlcN (from here on this condition is named Rieske-iKD and C3AP2-iKD respectively) and monitored the growth over four and seven 48h-long intra-erythrocytic cycles (IDCs) respectively. We did not observe a growth defect in C3AP2-iKD even after seven IDCs (Figure 4A). On the other hand, a severe growth defect at three IDC was seen upon PfRieske depletion (Rieske-iKD) (Figure 4B). Giemsa stained Rieske-iKD parasites seem morphologically similar to their parental control for two IDCs and the growth defect was observed at the late-trophozoite-to-early-schizont stage in the third IDC which is confirmed by stage quantification analysis of the Rieske-iKD parasites compared with the parental control (Figure 4C-F). These data indicate that the PfRieske subunit is essential for asexual parasites development.

**Figure 4.**
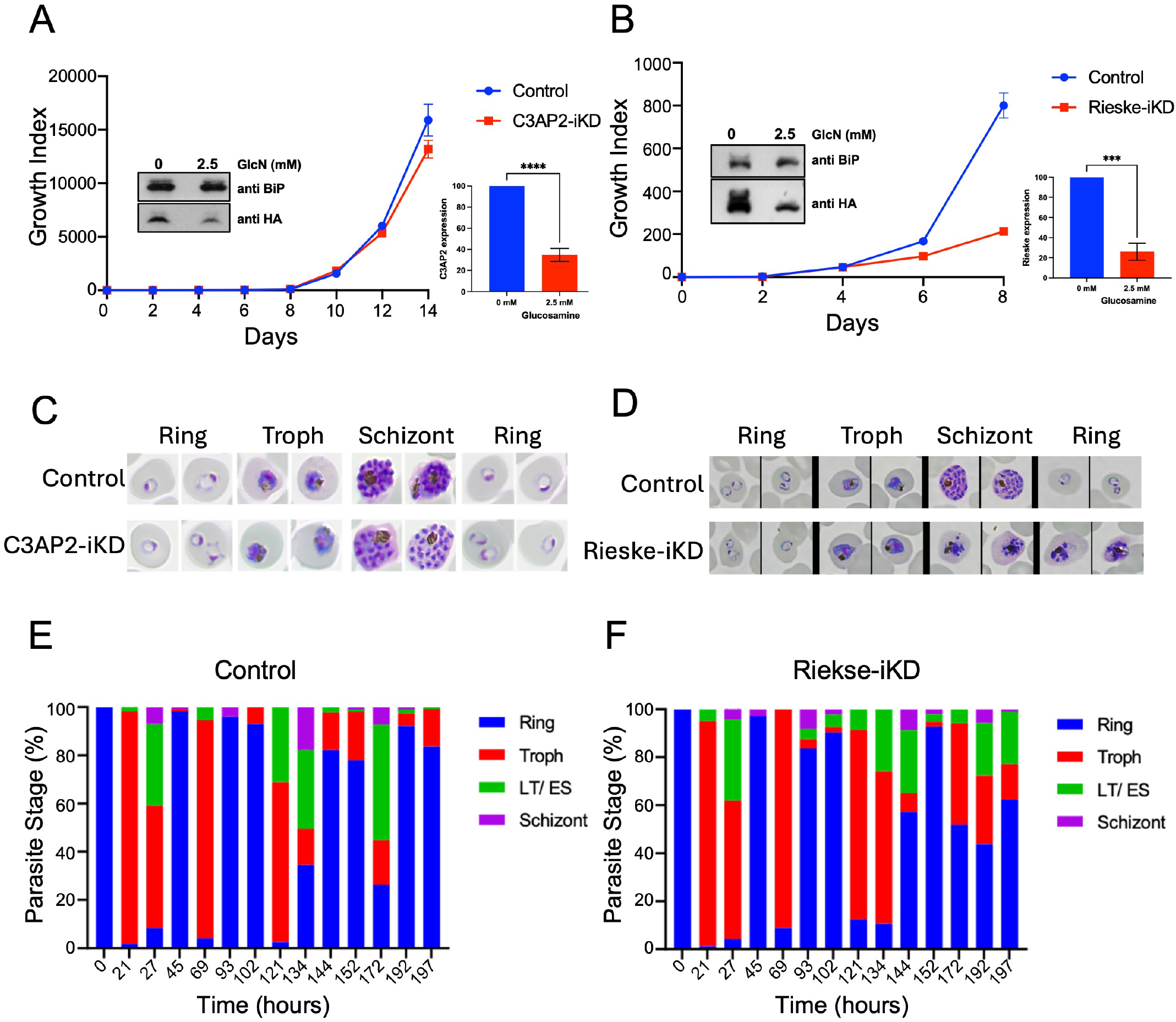
Inducible knockdown shows that PfRieske is essential for the asexual blood stage parasite growth: (A,B) Growth curves of C3AP2-iKD n = 2 (A) and Rieske-iKD n = 3 (B) transgenic parasites growing in presence and absence of the 2.5 mM glucosamine (GlcN). Insets are showing western blots probed with anti-HA and anti-BiP antibodies of the parasite cell lysate grown in GlcN presence and absence condition. Error bars are SEM. (C,D) Representative images of Giemsa-stained smears of the third cycle of the C3AP2-iKD (C) and Rieske-iKD (D) parasites grown in presence or absence of GlcN. (E,F) Bar graph showing quantification of the life stage progression of the Rieske-iKD parasites in absence (E) and presence (F) of GlcN n = 2 for both. LT/ES – late trophozoite/early schizonts.

### Loss of PfRieske results in hypersensitivity to mETC inhibitors and to membrane depolarisation by proguanil

Several compounds are well-established inhibitors of enzymes involved in the transport of electrons in the Plasmodium mitochondrion. PfCIII is known to be inhibited by atovaquone [8-10]; DHODH is targeted by DSM-265 [26]; and, while the direct target of proguanil is not fully resolved, it is believed to disrupt the mitochondrial membrane potential too [6, 27]. Parasites with decreased expression of the targets of those compounds are expected to be hypersensitivity to their inhibition. This was demonstrated in a study of the Plasmodium mitochondrial ribosome (mitoribosome). Disruption of the mitoribosome leads to a decrease in functional PfCIII because one of the PfCIII components, cytb, is encoded in the mitochondrial genome, and indeed such disruption led to hypersensitivity to atovaquone, DSM-265 and proguanil [28]. Likewise, as we suspect that PfRieske depletion in our line results in decrease of functional PfCIII, we hypothesised that it would lead to similar hypersensitivity. Seeing that the growth defect following PfRieske depletion is detected on day six (third IDC), we added inhibitors on day four (at the ring stage of the second IDC) and monitored the parasite growth over the next cycle, IC50 were then calculated on day six (Figure 5A). In line with our hypothesis, Rieske-iKD parasites show hypersensitivity towards atovaquone, DSM-265, and proguanil (Figure 5B, Table S2). The IC50 for chloroquine and artemisinin, which do not interfere with the mETC, remained unchanged compared to wild type (Figure 5C, Table S2), indicating that the observed hypersensitivity is specific to inhibition of this pathway.

**Figure 5.**
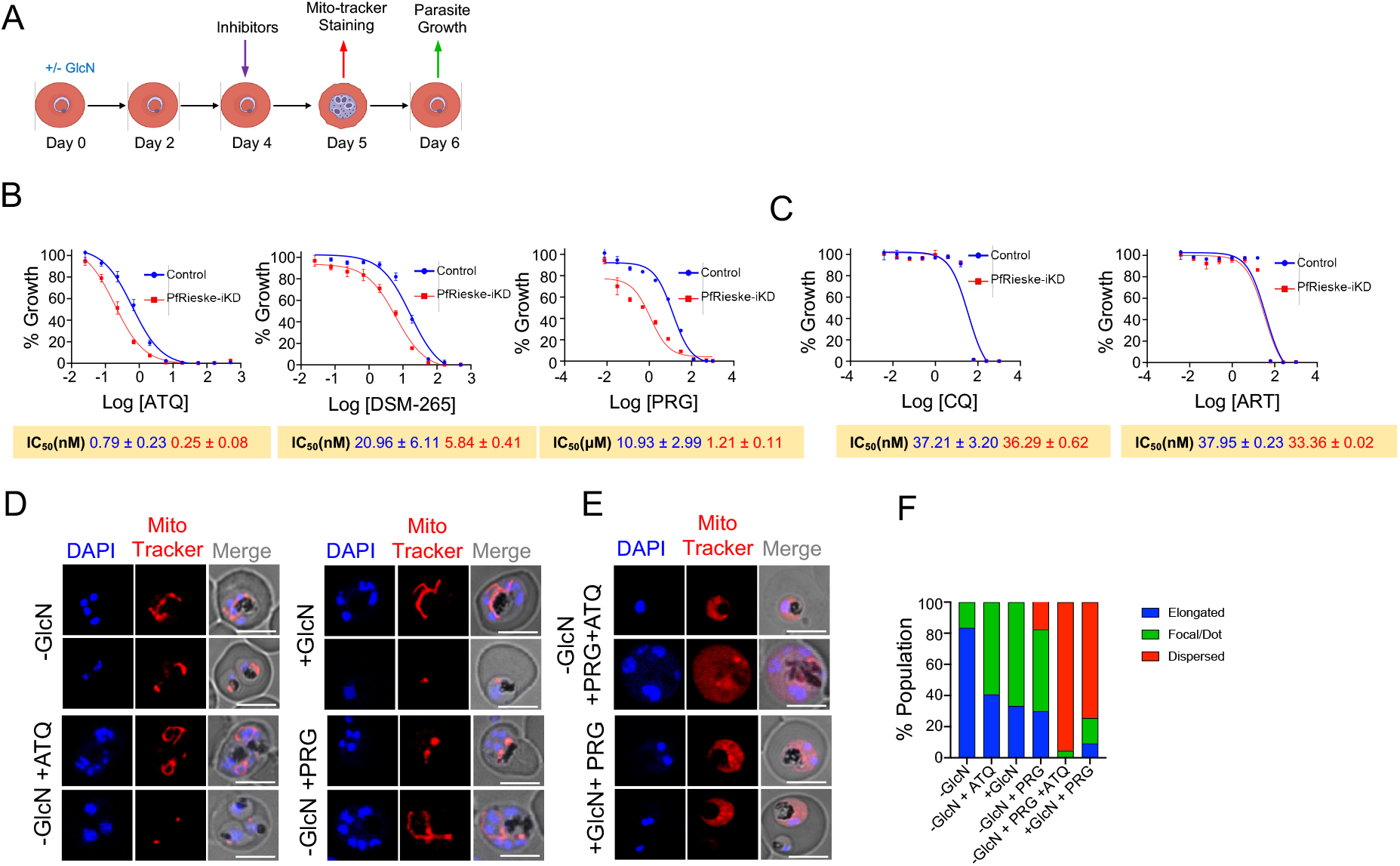
Depletion of PfRieske results in mETC failure: (A) Schematic representation of our experimental setup for the membrane potential and hypersensitivity assays. (B) IC50 analysis reveals hypersensitivity of GlcN treated parasites (Rieske-iKD, compared to the control which is the same parasite line not treated with GlcN) to the mETC inhibitors atovaquone (ATQ), DSM-265 and proguanil (PRG). (C) IC50 analysis confirms no change in the sensitivity of GlcN treated parasites (Rieske-iKD) to drugs used as negative control for (B), Chloroquine (CQ) and Artemisinin (ART). The graphs in B and C are each a representative from n=2 independent experiments, and the numbers are the mean of the technical triplicates. Table S2 provides the values from both experiments. (D,E) confocal fluorescent microscopy images of MitoTracker-Deep Red stained parasites grown in the presence or absence of glucosamine, atovaquone and proguanil (-/+ GlcN/ ATQ / PRG). Combinations that do not affect the membrane potential are shown in (D) as controls, while treatment with atovaquone and proguanil, or upon PfRieske depletion and proguanil treatment are shown in (E). Scale bar: 5μm (F) Quantification of the mitochondrial signal in each of the conditions in (D) and (E). The data shown is representative of the n=2 independent experiments, 50 parasites analyzed.

Since electron transport is coupled to proton pumping, which in Plasmodium is expected to occur in Complexes III and IV, we reasoned that PfRieske depletion will reduce the mitochondrial membrane potential. We used MitoTracker-Deep Red as a proxy for membrane potential and investigated the effect of PfRieske depletion and mETC inhibitors on its signal distribution. Rieske-iKD parasites grown in presence of GlcN for five days still presented MitoTracker-Deep Red mitochondrial specific staining, as did parasites treated with atovaquone (1 nM) or most parasites treated with proguanil (1 μM) alone (Figure 5D). However, nearly all parasites treated with both atovaquone and proguanil lose the membrane potential dependent mitochondrial staining of Mito-Tracker-Deep Red and the signal was detected in the cytosol instead (Figure 5E). Likewise, treatment of PfRieske-HA-glmS with both GlcN and proguanil resulted in loss of the mitochondrial staining (Figure 5E). These data indicate that the depletion of PfRieske disrupts the membrane potential in a manner that mimics the disruption caused by inhibition with 1 nM of atovaquone.

### PfRieske is required for the pyrimidine synthesis pathway

The profile of hypersensitivity to inhibitors and the observed disruption of the mitochondrial membrane potential upon PfRieske depletion is in line with a decrease in functional PfCIII in this mutant. Previous studies demonstrated that the main function of PfCIII in asexual stage parasites is in ubiquinone recycling for the DHODH-ubiquinone oxidoreduction cycle which is an essential step in the pyrimidine synthesis pathway [6]. We thus hypothesised that the observed growth defect is due to a defect in the pyrimidine synthesis pathway. Since we observed the growth arrest in the third cycle (Figure 4B) we isolated whole-cell metabolites from the trophozoite stage of this cycle (on day five) and performed targeted metabolomic analysis for intermediates of the pyrimidine synthesis pathway that are found upstream of DHODH: N-carbamoyl-L-aspartate and dihydroorotate (Figure 6A). We found that the level of both metabolites was significantly enhanced (∼4 folds increase) in the Rieske-iKD (Figure 6B), suggesting that they accumulate in this mutant. As a positive control we analysed those metabolites in parasites that were treated with 10 nM atovaquone for four hours which also resulted in the accumulation of these metabolites as expected (Figure 6B).

**Figure 6.**
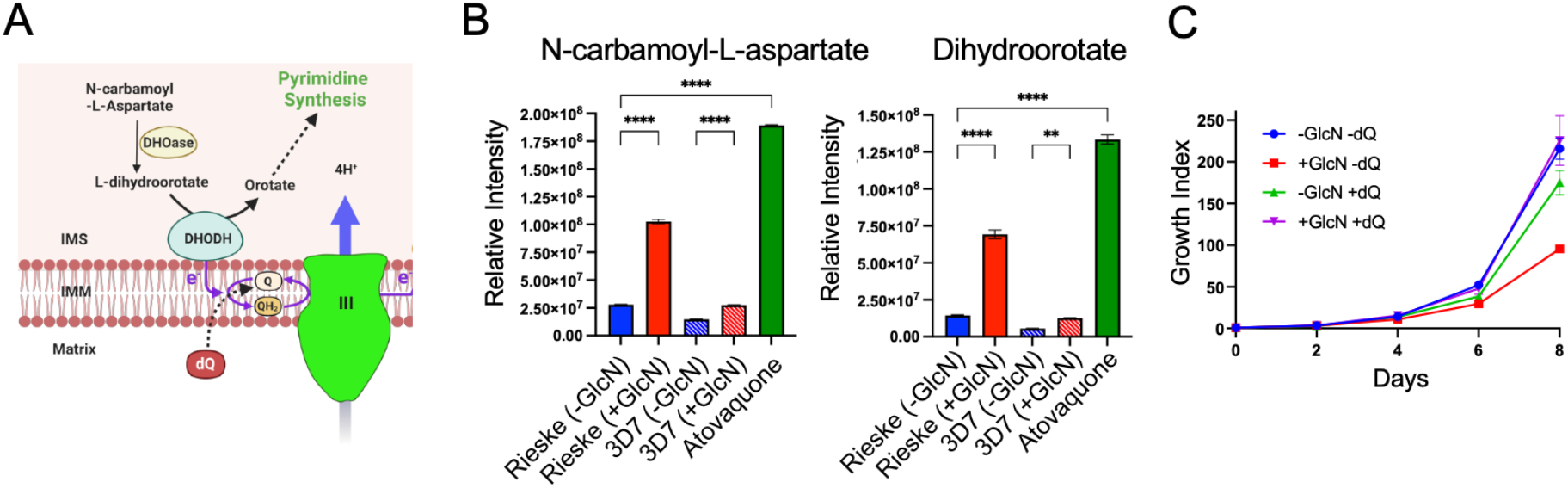
Accumulation of pyrimidine synthesis precursors in Rieske-iKD, while treatment with decyl-ubiquinone rescues its growth defect: (A) schematic image of the part of the pyrimidine synthesis pathways involving DHODH, ubiquinone and PfCIII. The mitochondrial intermembrane space (IMS), inner membrane (IMM), Ubiquinol (QH2), ubiquinone (Q) and decyl-ubiquinone (dQ) are shown. (B) Bar graph showing the levels of the analysed pyrimidine synthesis pathway intermediates, N-cabamoyl-L-aspartate (left) and Dihydroorotate (right), in parental and Rieske-iKD lines grown in the presence or absence of GlcN (Rieske / 3D7 (-/+ GlcN) or upon treatment with atovaquone. (C) Growth curve of Rieske-iKD in presence and absence of GlcN (-/+ GlcN) and of 30 μM dQ (-/+ dQ). Errors bars are SEM.

Previous studies have demonstrated that the need for PfCIII to oxidize ubiquinol to ubiquinone for the cycle of DHODH-ubiquinone oxidoreduction can be bypassed through addition of the ubiquinone analogue decyl-ubiquinone (dQ) to the parasite growth media [16, 29, 30]. It would therefore be expected that the observed growth defect upon PfRieske depletion should be rescued with dQ. Accordingly, parasites grown in the presence of both GlcN and dQ showed restored growth comparable to parasites grown in the absence of GlcN (Figure 6C). Taken together these data provide additional support that PfRieske is required for PfCIII function and suggests that the observed growth defect of asexual stages in our mutant is likely due to a defect in pyrimidine synthesis.

### Loss of PfRieske causes decrease in cristae density and affects gametocytes maturation

The transition from asexual stages to gametocyte is accompanied by a shift in carbon metabolism whereby gametocytes rely more on the TCA cycle, mETC and ATP synthase for energy and metabolites [31, 32]. In line with this shift, mETC components show up to 40-fold increase in expression in the sexual stages [15]. We therefore expect that PfCIII will play a critical role in more pathways in gametocyte biology compared to the asexual stages. This is supported by the finding that development and transmission in mosquitos depends on a fully functional PfCIII [21, 33, 34]. We were therefore interested to examine the impact of PfRieske depletion on gametocytes development. We induced gametocyte commitment using the spent-medium protocol [35] (Figure 7A,B). Morphological analysis revealed that gametocyte development was severely affected upon PfRieske depletion: in the control set, 100 % of the parasites analysed reach stages IV/V, whereas ∼30% abnormal gametocytes were seen in Rieske-iKD (Figure 7B,C). Abnormal gametocytes were characterised with round overall morphology and what seem to be a swollen food vacuole (Figure 7B,D).

**Figure 7.**
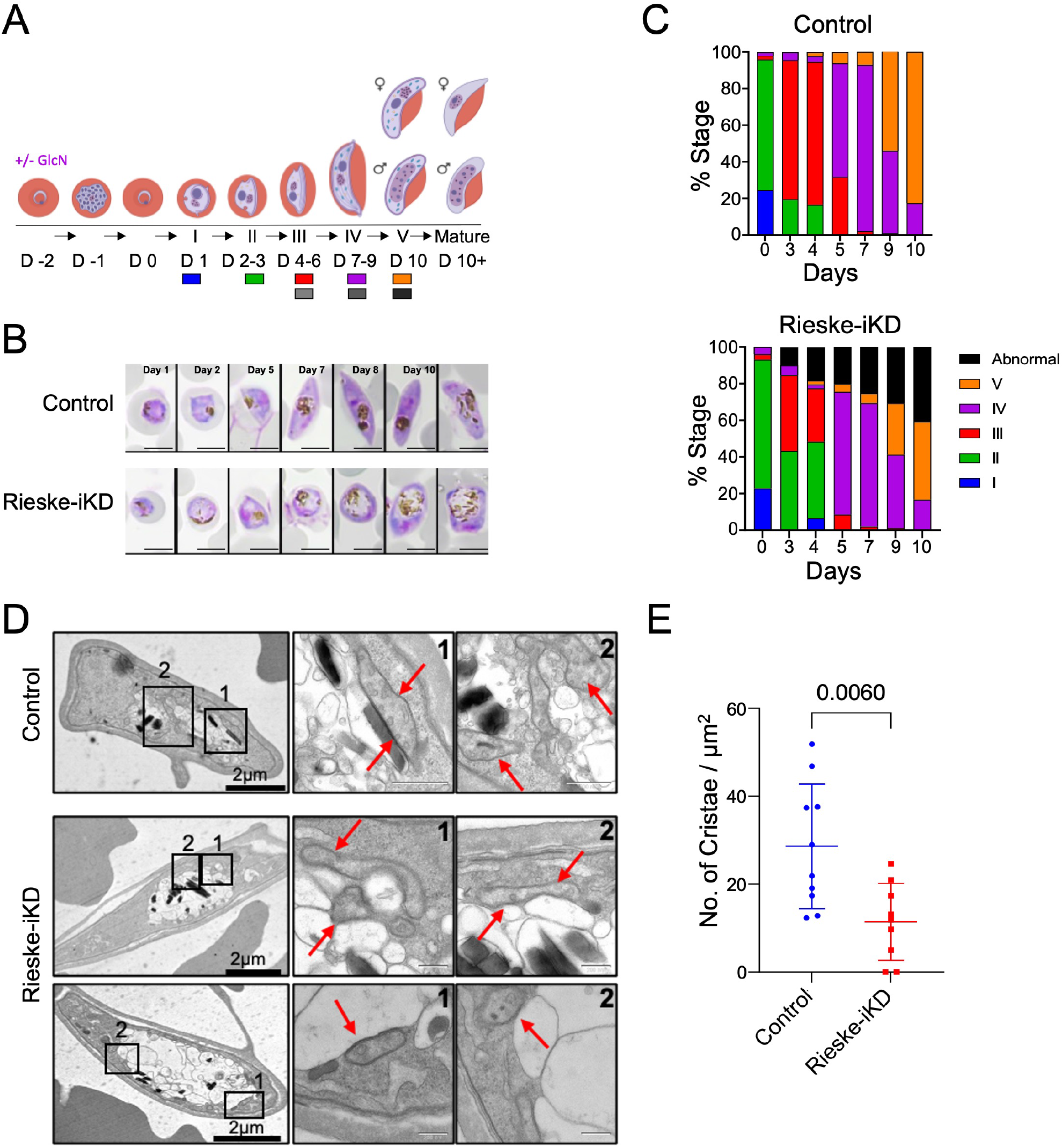
Knockdown of PfRieske arrests gametocyte development: (A) Schematics of the experimental setup of gametocyte induction in control and Rieske-iKD along with a scheme of the different stages. (B) Progression of the control and Rieske-iKD gametocytes monitored by giemsa stained smears. Scale bar: 5μm (C) quantification of morphologies based on giemsa-stained smears of control and Rieske-iKD gametocytes. (D) Transmission electron micrographs of the control and Rieske-iKD gametocytes. Enlarged sections show mitochondria (red arrows). Scale bar: 200nm (E) Quantification of cristae density in the control and Rieske-iKD gametocytes.

Mitochondrial cristae shape is tightly linked to oxidative phosphorylation (OXPHOS) activity. Accordingly, the metabolic shift in gametocytes coincides with de novo appearance of cristae [7, 36, 37]. We thus hypothesised that the defect in gametocyte development upon PfRieske depletion would be linked to cristae abnormalities. In support of this hypothesis, examination of the Rieske-iKD parasites using transmission EM (TEM) revealed abnormal mitochondrial morphology upon PfRieske depletion (Figure 7D). Quantification of cristae numbers per mitochondrial area further showed that these parasites have a reduced cristae density in their mitochondria compared with their parental line (Figure 7E). To our knowledge this is the first time that a cristae defect is observed upon disruption of a Plasmodium gene. These data show that PfRieske is required for maintaining the cristae density in *P. falciparum* gametocytes which may ultimately help the parasite in fulfilling the metabolic needs required for gametocyte maturation.

## Discussion

The mETC is critical for the survival of many eukaryotic cells including both the apicomplexans and their hosts, nevertheless the parasite pathway shows different sensitivity to inhibitors compared to the human one. For this reason, the *Plasmodium* mETC has been the focus of numerous drug discovery studies for antimalarial drug development, including a recent screen of the Medicines for Malaria Venture (MMV) “pathogen box” [38]. Most of our knowledge about *Plasmodium* mETC complexes including PfCIII is therefore based on studies using inhibitors or analysing drug-resistance, while their role within parasite biology is less well understood. Here we use genetic disruption to provide improved understanding of the role of PfCIII in parasite biology in asexual and sexual stages.

Recent studies in apicomplexans proposed a divergent set of mETC complex subunits compared to the mETC of the host and highlighted new parasites subunits in each of the complexes [15, 17, 39, 40]. PfCIII was proposed to consist of 12 subunits with a predicted higher molecular size compared to human complex III [15]. Via C-terminal tagging we provided support for this prediction by recovering some of the new subunits in immunoprecipitation and via BN-PAGE and western analysis which confirmed that both the conserved PfRieske and the new subunit PfC3AP2 are part of a complex which migrates at size of ∼730 kDa. It will be of interest to understand how the new subunit contribute to PfCIII function and what is their mechanism of action in this important complex. One possibility is that they would structurally “fill-in” for the missing homologs of the non-catalytic subunits found in other systems, as we recently proposed for the non-catalytic subunits of mETC complex II in the related *T. gondii* parasite [40]. Another option is that they may mediate interaction with other mETC complexes to form supercomplexes, as suggested for non-catalytic and species-specific subunits in other systems [41]. Importantly, regardless of their function, the presence of different proteins in PfCIII compared to its human counterpart may well induce small changes in the location and orientation of amino acid in and around the substrate binding pockets that could affect the interaction of PfCIII with inhibitors and that may provide opportunities for new drug development. Structural analysis of PfCIII will likely help shed light on these open questions.

Considering the essential role of PfCIII demonstrated in previous inhibitor studies, it is expected that its subunits will be important for parasite survival. In agreement with this expectation a previous study shows that a conditional depletion of mACP which leads to secondary loss of PfRieske ultimately leads to asexual stage parasite death [30]. Likewise, depletion of cytC1 protein demonstrated its essentiality for mETC functions and growth of asexual stages [16]. Our conditional knockdown adds support to these observations by showing directly that PfRieske is essential for asexual growth. However, to our surprise, PfC3AP2 protein downregulation did not have a significant impact on growth. This was unexpected given that in the genome wide piggy bac transposons-based study PfC3AP2 is predicted to be essential for growth [42], and our previous study of the PfC3AP2 homolog from *T. gondii* also showed that it is essential for parasite survival [17]. We cannot fully conclude weather the regulation accomplished with the glmS system is not sufficiently tight or weather PfC3AP2 is truly dispensable for asexual blood stage parasites at this point.

Previous studies demonstrated that disruption of a component of PfCIII [16] and of the *P. falciparum* mitoribosome [43] (required to translate the essential PfCIII subunit cytb), leads to parasites hypersensitive for mETC inhibitors, likely since there is less PfCIII activity remaining to inhibit in those mutants. Studies further suggested that a potential alternate pathway to polarise the inner mitochondria membrane independent of complex III and complex IV exists in *Plasmodium* [6, 44]. This pathway can be blocked by proguanil only in the presence of a complex III inhibitor. As reported upon depletion of PfCIII component, cytC1 [16], we observed that loss of PfRieske alone did not result in membrane depolarisation, which instead required a combination of PfRieske depletion and treatment with proguanil. Collectively, our observations thus agree with previous findings, thus providing support that PfRieske depletion results in PfCIII disruption in our case too.

The mETC plays two major cellular functions: the one is the electron transfer, whereby the transported electrons reduce a final electron acceptor, often oxygen. This electron transfer function enables different metabolic pathways through oxidation of key enzymes, such as DHODH of the pyrimidine synthesis pathway; the other is oxidative phosphorylation, whereby the electrons transported from different metabolic pathways are coupled to the pumping of protons by mETC complexes, generating an electrochemical gradient that facilitates ATP synthesis by ATP synthase. The *P. falciparum* life cycle is complex, consisting of multiple developmental stages and requiring survival in two hosts. The different stages and the moves between host environments manifest in major metabolic shifts and the reliance on different roles of the mETC is expected to vary accordingly. For example, it was established that in the asexual blood stages the main function of *P. falciparum* mETC is enabling the oxidation of DHODH for *de novo* pyrimidine synthesis [6]. Our metabolomic data confirms that PfRieske depletion results in defect in the pyrimidine synthesis pathway and accumulation of two key metabolites upstream of DHODH. Likewise, the ability of dQ to rescue the growth of PfRieske depleted parasites, which is in line with previous observations [16, 29], provides further support for the sole role of ubiquinone recycling that PfCIII plays in the asexual blood-stages.

On the other hand, different findings point to the importance of active oxidative phosphorylation in the transmissible stages, including the increased expression of genes encoding mETC components in gametocyte [15], and the growth defect of the mosquito oocysts stages upon disruption of other oxidative phosphorylation enzymes such as complex II, ATP synthase and NDH2 in *P. berghei* [7, 45, 46]. Likewise, mitochondrial cristae, which are a hallmark of active oxidative phosphorylation are almost only present in the sexual stage parasites. In line with these observations, we found a defect in gametocyte development upon PfRieske depletion, and this outcome coincides with reduced cristae density in those PfRieske depleted gametocytes. In another protozoan, *Tetrahymena thermophila*, complex III is a part of a supercomplex that plays a role in curving the cristae [47]. It is possible that one of the PfCIII containing supercomplexes that were detected in *Plasmodium* and that were found to be more abundant in sexual compared with asexual stages [15] plays a similar role in these parasites. In this scenario, PfCIII depletion would lead to a decrease in the number of any supercomplexes involving it, and thus to the observed cristae formation defect. However, the mechanism controlling cristae biogenesis in *P. falciparum* requires further studies at this point. Another possibility is that the reduced level of PfCIII is not sufficient to support oxidative phosphorylation and that the change of cristae morphology occurs in respond to a reduced rate of respiration. This is in line with previous evidence of the critical role of PfCIII in gametocyte metabolism showing that atovaquone treatment affects gametocyte maturation and viability [48]. Finally, we cannot exclude a scenario whereby the effect seen in gametocyte is secondary to the pyrimidine synthesis defect observed in the asexual stages.

Interestingly, on top of the mitochondrial ultrastructure defect, PfRieske depleted gametocytes also show an enlarged food vacuole. Seeing that the food vacuole plays a central role in parasite metabolism via haemoglobin degradation and recycling of amino acids, we speculate that the functions of these two organelles are tightly coordinated and that the two phenotypes are potentially linked. The apicomplexan mitochondrion is seen to be involved in active membrane contact sites with several organelles (ER, pellicles, nucleus [49-53]). It would be of interest to explore if there are contacts with the food vacuole too.

## Materials and Methods

### Plasmid construction, parasite culture, transfection, and transgenic line confirmation

To generate the HA-glmS construct, C-terminal region of PfRieske and PfC3AP2 were PCR amplified using primer sets (2448,2449 for C3AP2 and 2451, 2535 for Rieske) and cloned into HA-glmS vector in *BglII* and *PstI* restriction sites. *Plasmodium falciparum* 3D7 strain was cultured under standard culture conditions in RPMI media supplemented with 0.5% (W/V) Albumax (Invitrogen). To generate the HA-glmS transgenic parasite lines, 3D7 parasites were synchronised repeatedly by sorbitol treatment and 100 ug of plasmid DNA for the respective gene was transfected in *P. falciparum* by electroporation (0.310 kV and 950 μF) [53]. Transfected parasites were selected with WR 99210 (Jacobus Pharmaceuticals). Clonal selection was done by serial dilution in 96 well plate and the integration were confirmed by PCR using primer set (2718, 2797 for C3AP2 and 2719, 2797 for Rieske) and by western blot analysis using anti HA antibody.

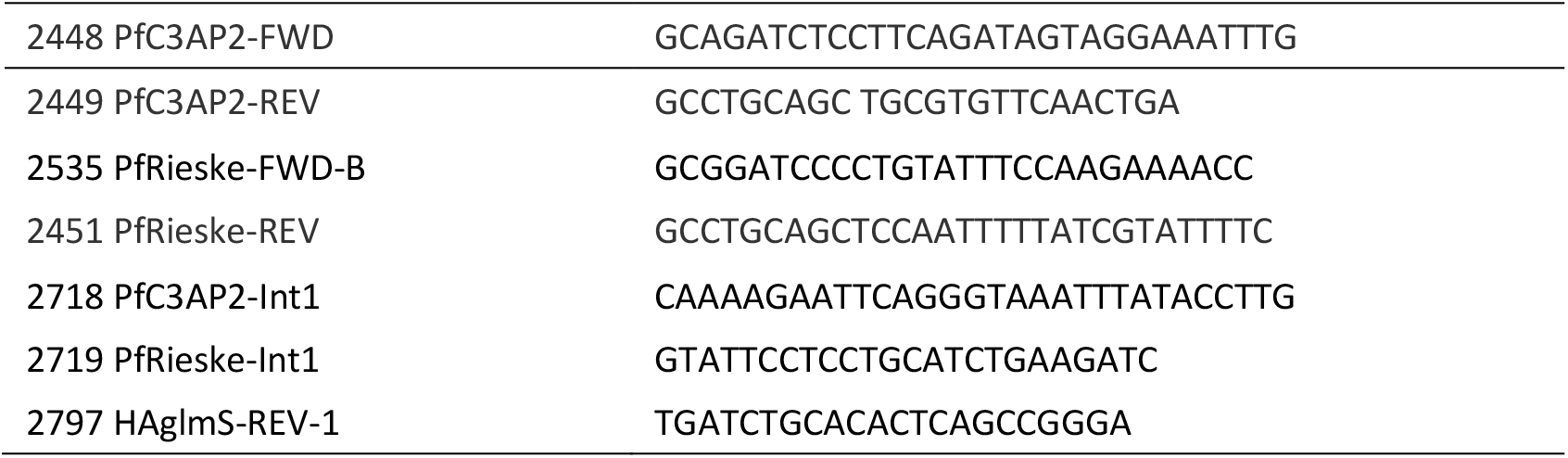

### Inducible knockdown and parasite growth assay

Both the transgenic parasite lines were tightly synchronized with repeated 5% sorbitol treatment. To assess the knockdown at the protein level, 2.5mM glucosamine (GlcN) was added at the ring stage and schizont stage parasites were harvested using saponin. The protein samples for western blotting were prepared from these parasites.

For growth assays, parasites were cultured in with/without GlcN conditions for several cycles and the parasitemia was assessed at different intervals by malaria SYBR Green I-based fluorescence assay (MSF) using MACSQuant. Briefly, cells were incubated with SYBR green for 20 minutes at 37 °C in dark followed by the 3 washings with 1 X PBS and these samples were analysed MACSQuant system. For each set 100000 cells were counted to monitor the parasitemia. Uninfected RBCs were used as negative control. The growth index is the cumulative parasitemia, which is the multiplication of parasitemia and split factors (1:4 split every IDC) over the time course.

To assess the stage specific knockdown effect, giemsa stained smears were prepared at different interval from the control and GlcN treated parasites and analysed by light microscope. Thousand infected RBCs (iRBCs) were counted to check the stage specific defect of the Rieske-iKD.

### Mitochondria enrichment and Immunoprecipitation

Both the transgenic parasite lines and the parent 3D7 parasites were grown at large scale under standard cultured conditions. For each line, eight large dishes of 10% late trophozoite-schizont stage parasites were harvested using the 0.15% saponin treatment. Parasites pellets were washed with 1X PBS several times to remove the haemoglobin contamination. Enriched mitochondrial fraction was purified from these parasites using the previous published protocols. Briefly, saponin purified parasites cells were broken using the nitrogen cavitation at 1500 p.s.i. for 20 minutes and passed through the miltenyi Biotech MACS column to remove the hemozoin. This material was further pelleted by centrifugation and used for further experiments.

### Immunoprecipitation and mass spectrometry

The mitochondrial samples were resuspended into the lysis buffer (1 % DDM) and kept on rotor in cold room for 2 hours for lysis. These were centrifuged at 16000 x g for 30 minutes and the supernatant was used for the overnight binding on the Pierce anti-HA agarose beads. Next day, beads were washed thrice with the lysis buffer and proteins were eluted by incubating the beads with HA peptide (1mg/ml) overnight. Samples were collected at different steps and analysed by western blotting of both SDS-PAGE and BN-PAGE. Beads and eluted samples were sent for mass spectrometry analysis.

For Mass spectrometry analysis trypsin digested peptide samples were prepared and these dry peptides residues were solubilized in 20 μL 5 % acetonitrile with 0.5 % formic acid using the auto-sampler of a nanoflow uHPLC system (Thermo Scientific RSLC-nano). Online detection of peptide ions was by electrospray ionisation (ESI) mass spectrometry MS/MS with an Orbitrap Elite MS (Thermo Scientific). The ionisation of LC eluent was performed by interfacing the Sharp Singularity emitters (Fossil Ion Tech) with an electrospray voltage of 2.5 kV.

An injection volume of 5 μL of the reconstituted protein digest were desalted and concentrated for 1.1 min on trap column (0.3 × 5 mm) using a flow rate of 25 μl / min with 1 % acetonitrile with 0.1 % formic acid. Peptide separation was performed on a Pepmap C18 reversed phase column (50 cm × 75 μm, particle size 3 μm, pore size 100 Å, Thermo Scientific) using a solvent gradient at a fixed solvent flow rate of 0.3 μl / min for the analytical column. The solvent composition was A) 0.1 % formic acid in water B) 0.08 % formic acid in 80% acetonitrile 20% water. The solvent gradient was 4 % B for 1.5 min, 4 to 60 % for 100.5 min, 60 to 99 % for 14 min, held at 99 % for 5 min. A further 9 minutes at initial conditions for column re-equilibration was used before the next injection.

Protein identifications were assigned using the Mascot search engine (v2.6.2, Matrix Science) to interrogate protein sequences in the *Plasmodium falciparum* 3D7 database. A mass tolerance of 10 ppm was allowed for the precursor and 0.3 Da for MS/MS matching.

### Denaturing polyacrylamide gel electrophoresis (SDS-PAGE), Blue native (BN-PAGE), and western blotting

For SDS-PAGE, Parasites were isolated by lysing the infected RBCs with 0.15% saponin and the parasites were resuspended in Laemmli buffer and lysed by heating at 95 for 10 minutes. Proteins were further separated on the 12% SDS-PAGE and transferred on the PVDF membrane (Millipore) using semi dry trans-blot (Bio-Rad). For the BN-PAGE, samples were prepared by resuspended the protein samples in solubilisation buffer and incubated on ice for 30 minutes and centrifuged at 16000 x g at 4C for 30 minutes. The supernatant was combined with sample buffer containing Coomassie G250 (NativePAGE) at the final concentration of 0.25% DDM and 0.0625% Coomassie G250. The samples were separated on the 4-16 % Bis-Tris native gel and transferred on PVDF membrane.

For the western blotting, membranes were incubated with blocking solution (5% skimmed milk in 1X PBS) followed by overnight incubation with respective primary antibodies (anti HA rat 1:1000 (Roche), anti BiP rabbit 1:5000 in cold room and 1 hr incubation with respective secondary antibodies (1:10000) at room temperature and visualised by using ECL kit (Thermo-scientific).

### Immuno-fluorescence assay (IFA) and microscopy

Infected RBCs (5-10% parasitemia) were fixed in 4% paraformaldehyde and 0.05% glutaraldehyde for 50 min at room temperature on rotator. Cells were permeabilised by 10 minutes incubation with 0.1% Triton X-100 on ice and incubated with blocking buffer (10% FBS in 1X PBS) for 90 minutes and then probed with the primary antibody (anti-HA rat 1:100 (Roche) and antiPfs25 rabbit 1:1000) for 2 hours on rotor and after three washing with 1x PBS cells were probed with anti-rat Alex Flour-488 (1:500) for 1 hour. Parasites nuclei were stained with DAPI with a final concentration of 5ug/ml. Images were captured by Nikon A1 confocal laser scanning microscope and analysed by Nikon-NIS element software (version 4.1).

To label mitochondria infected RBCs were incubated with 100nM Mito Tracker Deep Red for 15 minutes at 37 °C shaker incubator and washed with 1x PBS thrice. Cells were further fixed and processed for IFA as describes earlier.

### Metabolomic profiling

Rieske-HA-glmS parasites were synchronised, and knockdown assay was setup as describes earlier. Parasites were isolated from the control and Rieske-iKD parasites using saponin and 3×10^8^ number of parasites were used for metabolites isolation. Isolated parasites were quenched by rapidly cooling the cells to 4 °C by dry ice/ethanol. Metabolites were isolated using chloroform/methanol/water (1:3:1) method. Briefly, cells were resuspended in 600 μL of chloroform/methanol/water (1:3:1) and kept on vortex for 1 hr at 4 °C. After centrifugation for 3 minutes at 13000 x g at 4 °C, supernatant was transferred in to new eppendorf tube and stored in −80 °C until analysis by LC-MS.

Hydrophilic interaction liquid chromatography (HILIC) was carried out on a Dionex UltiMate 3000 RSLC system (Thermo Fisher Scientific, Hemel Hempstead, UK) using a ZIC-pHILIC column (150 mm × 4.6 mm, 5 μm column, Merck Sequant). The injection volume was 10μl and samples were maintained at 5°C prior to injection. For the MS analysis, a Thermo Orbitrap QExactive (Thermo Fisher Scientific) was used.

Metabolomics samples from the control samples (3D7 -/+ GlcN, 3D7 treated with ATQ) also prepared and analysed in the same way.

### Drug sensitivity assay

The drug assay was performed in 96 well plate. Rieske-HA parasites were grown in media with/without GlcN for two cycles and second cycle ring parasites were exposed to drugs diluted in serial dilution. In the third cycle parasites were lysed in SYBR green lysis buffer and fluorescence intensity was assayed using spectrophotometer. IC_50_ graph for drugs were prepared using Graph prism software. All the compounds used in this assay were purchased Atovaquone, DSM-265, Proguanil, Chloroquine, Artemisinin

### Transmission electron microscopy

For electron microscopy analysis, control and Rieske-iKD gametocytes were fixed with prewarmed 2% glutaraldehyde in 0.1M cacodylate (pH 7.4) buffer. Following serial washes in 0.1M phosphate buffer, samples were post-fixed in 1% OsO4 and 1.25% potassium ferrocyanide (vol:vol) in the same buffer for 1 hour in the dark, washed with distilled water and contrasted EM bloc with 0.5% aqueous uranyl acetate for 1 hour at room temperature in the dark. The samples were then dehydrated in acetone ascending series (30%, 50%, 70%, 90%, 100%) and embedded in epoxy resin. Ultra-thin sections (50nm) were sectioned in a Leica Ultramicrotome and collected in 100mesh grids covered with formvar. The collected sections were contrasted with 2% aqueous uranyl acetate and later observed in a Jeol 1200 transmission electron microscope (Jeol, Japan) operating at 80kV.

### Gametocyte commitment and culturing

Gametocyte commitment assay was induced by stress using spent media. Briefly, Rieske-HA-glmS parasites were synchronised and divided in GlcN (+) or GlcN (-) conditions one cycle before the commitment. Gametogenesis commitment was induced using nutritional stress by grown parasites with spent medium (day two) and then growing parasites in media having heparin (1:500) till day +three to block the invasion of the asexual stage parasites. After that parasites were cultured with normal media conditions. Giemsa-stained smears were prepared at different time intervals for the morphological analysis.

## Supporting information

Supplementary Tables 1 ans 2

## Supplementary Materials

The following supporting information are added.

Table S1 – full list of the outcome from the proteomics analysis of PfRieske-HA immunoprecipitation. In yellow are subunits predicted to be part of PfCIII according to complexome studies and homology searchers.

Table S2 –Separate experiment results and calculation for the IC50 shown in Figure 5. Subunits predicted to be part of PfCIII according to complexome studies and homology searchers are shown.

## Author Contributions

Conceptualization, LS.; methodology, LS, PS.; validation, PS; formal analysis, PS.; investigation, PS.; resources, LS.; writing—original draft preparation, PS.; writing—review and editing, LS, PS, AM.; supervision, LS.; funding acquisition, LS, AM. All authors have read and agreed to the published version of the manuscript.

## Acknowledgments

We thank Dr. katarzyna modrzynska and Dr. Andrew Maclean for critical review of our manuscript. We acknowledge the support of Mathias Marti and his team with access to CAT3 facilities, and thank Gillian Parker for managing the facilities. We thank Leandro Lemgruber of the Glasgow Imaging Facility for his support and assistance in producing TEM data for this work. We thank Glasgow polyomics facility for their assistance in producing proteomics and metabolomics data for the work. We thank Prof. Paul Gilson (Burnet Institute) for sharing plasmids.

## Disclaimer/Publisher’s Note

The statements, opinions and data contained in all publications are solely those of the individual author(s) and contributor(s) and not of MDPI and/or the editor(s). MDPI and/or the editor(s) disclaim responsibility for any injury to people or property resulting from any ideas, methods, instructions or products referred to in the content.

